# Effective Combination Immunotherapy using Oncolytic Viruses to Deliver CAR Targets to Solid Tumors

**DOI:** 10.1101/843052

**Authors:** Anthony K. Park, Yuman Fong, Nanhai G. Chen, Brook Jeang, Dileshni Tilakawardane, John P. Murad, Sang-In Kim, Jianming Lu, Sandra H. Thomas, Stephen J. Forman, Saul J. Priceman

## Abstract

Chimeric antigen receptor (CAR)-engineered T cell therapy for solid tumors is limited by the lack of tumor-restricted and homogeneous expression of tumor antigens^1,2^. Therefore, we engineered an oncolytic virus to express a non-signaling, truncated CD19 (CD19t) protein for tumor-selective delivery, enabling targeting by CD19-specific CAR T cells. Infecting tumor cells with a chimeric oncolytic vaccinia virus coding for CD19t (OV19t) produced *de novo* CD19 surface-antigen expression prior to virus-mediated tumor lysis. Co-cultured CD19-CAR T cells secreted cytokines and elicited potent cytolytic activity against infected tumors. Using multiple mouse tumor models, intratumoral delivery of OV19t induced tumor expression of CD19t and improved tumor control following CD19-CAR T cell administration. CAR T cell–mediated tumor killing also promoted release of virus from dying tumor cells, which propogated tumor expression of CD19t. These data demonstrate a novel immunotherapy approach utilizing oncolytic viruses to promote *de novo* CAR T cell targeting of solid tumors.

**One Sentence Summary:** We describe a novel and effective combination immunotherapy utilizing oncolytic viruses to deliver de novo cell surface expression of CD19 antigen promoting CD19-CAR T cell anti-tumor responses against solid tumors.

## Introduction

A major challenge for CAR T cell therapy is the identification of antigens with truly tumor-restricted expression, which poses considerable safety concerns and potentially narrows the therapeutic window for solid tumors^3^. CD19 has been an ideal target for CAR T cells against hematological malignancies for several reasons, including its highly restricted expression on B-cells and acceptable off-tumor on-target properties^4^. Extensive studies using CD19-directed CAR T cells have recently resulted in landmark FDA approvals of CAR T therapies for patients with B-cell malignancies^5,6^. In addition to the shared expression of solid tumor antigens on normal tissue, most of these antigens also have heterogeneous expression, limiting the potential for effective and durable anti-tumor responses^3^. Many solid tumors, including triple-negative breast cancers and liver cancers, lack amenable tumor antigens for CAR T cell development. Therefore, novel approaches to introduce validated targets to tumor cells could potentially broaden the applicability of CAR T cell therapy for otherwise intractable solid tumors.

Oncolytic viruses (OV) are a promising treatment modality for solid tumors, largely because of their tumor-selectivity, desirable immunogenic properties, and ability to incorporate transgenes into their genome for targeted delivery to tumors^7^. OV have gained significant momentum in recent years due to their immune-stimulating effects both systemically and in the local tumor microenvironment. The first clinically approved OV, Talimogene laherparepvec (TVEC), is a genetically modified type I herpes simplex virus that expresses granulocyte-macrophage colony-stimulating factor (GM-CSF)^8^. In addition to, and in some respects a consequence of, the selective tumor cell replication and direct tumor cell lysis, TVEC initiates exposure of soluble tumor antigens and induction of host anti-tumor immunity. Recent combination approaches have exploited OV to bolster adoptive cellular immunotherapy and immune checkpoint inhibitors. Additionally, novel versions of OV have been genetically engineered to express cytokines to further improve tumor recruitment of T cells and immune checkpoint inhibitors, thereby enhancing overall anti-tumor immunity^9^.

Here, we have exploited the transgene delivery potential of OV to selectively infect and drive tumor-specific expression of a proof-of-concept CAR-targetable tumor antigen, a truncated non-signaling variant of human CD19 (CD19t). We have demonstrated robust CD19t expression on multiple tumor types infected with OV carrying the CD19t gene (OV19t), which promoted activation and tumor killing by CD19-specific CAR T cells. Using human tumor xenograft models as well as an immunocompetent mouse tumor model, we showed potent anti-tumor responses combining OV19t and CD19-CAR T cells. Importantly, CD19-CAR T cells promoted the early release of intact virus from dying tumor cells, which effectively amplified the combination therapy.

## Results

### OV effectively deliver CD19t to solid tumor cells in vitro

For these studies, we utilized a chimeric orthopox OV, designed to infect a large spectrum of tumors^10^, carrying CD19t under the control of the vaccinia synthetic early promoter (P_SE_) (Fig. 1A) that allows for immediate surface expression of CD19t prior to oncolytic viral-mediated tumor lysis. The hCD19t expression cassette was inserted into the J2R locus encoding thymidine kinase (*tk*)^10^. This modified OV is termed CF33-(SE)hCD19t, and for the remainder of these studies referred to as OV19t. Of note, the inserted hCD19t gene replacing *tk* did not significantly impact *in vitro* infection efficiency or killing activity against human tumor cells (Fig. S1, A and B).

**Fig. 1.**
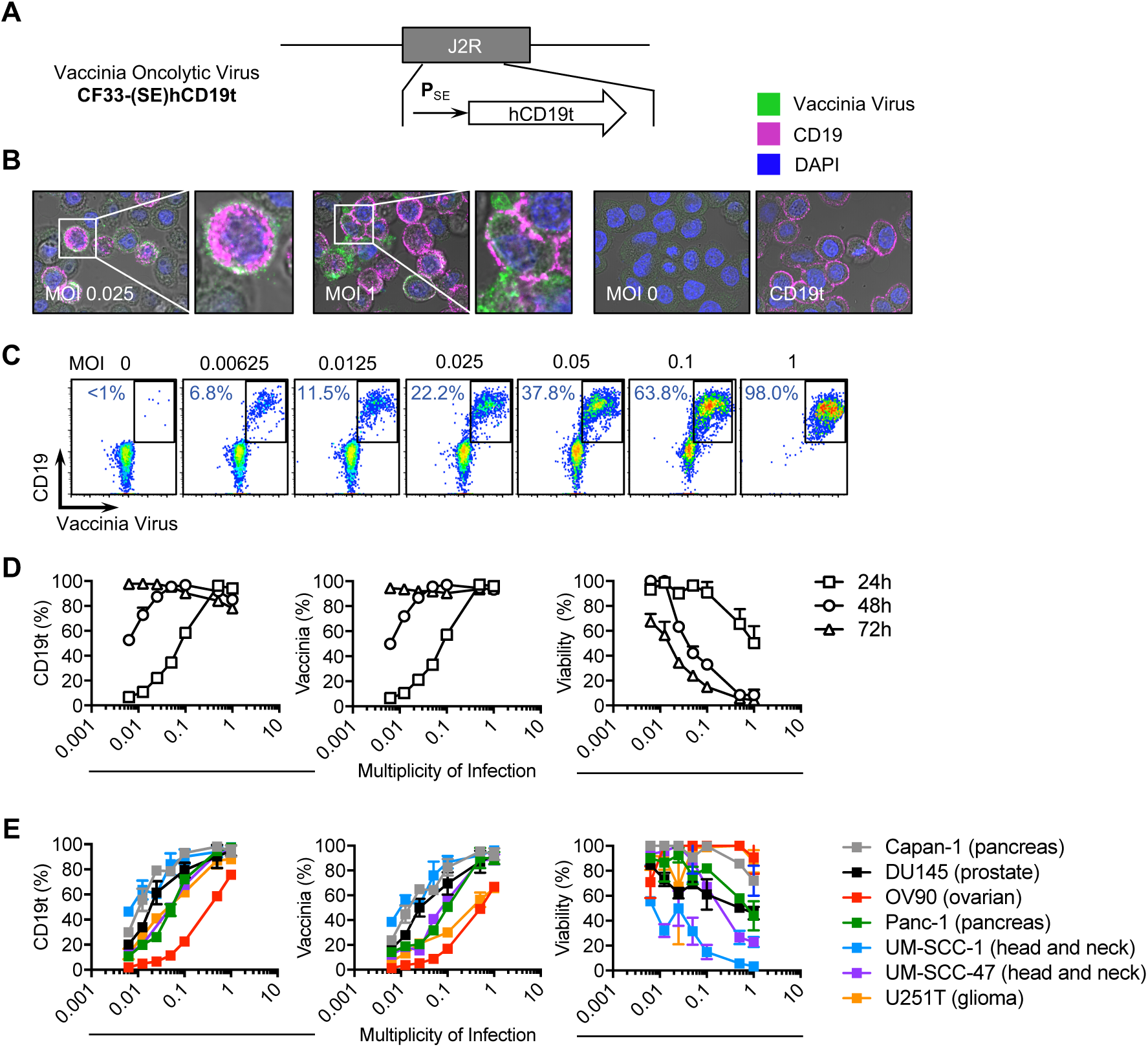
OV effectively deliver CD19t to solid tumors *in vitro*. A) Schematic of vaccinia OV [CF33-(SE)hCD19t], showing incorporation of human truncated CD19 (CD19t) under the control of the synthetic early promoter (P_SE_) inserted into the J2R locus replacing the thymidine kinase gene. B) Immunofluorescence microscopy of MDA-MB-468 cells infected for 24 h with OV19t at MOI 0.025 (left) and MOI 1 (middle), untransduced (MOI 0), or cells transduced with lentivirus to stably express CD19t (right). Images are at 10x magnification, and insets are at 40x magnification. C) FACS plots displaying the cell surface expression of CD19t and intracellular expression of vaccinia virus in MDA-MB-468 tumor cells determined by flow cytometry after 24 h of OV19t infection at increasing MOIs. D) Quantification of CD19t (left), vaccinia (middle), and viability (right) of MDA-MB-468 tumor cells following 24, 48, and 72 h co-culture with indicated MOIs of OV19t. E) Quantification of CD19t (left), vaccinia (middle), and viability (right) of indicated solid tumor cell lines following 24, 48, and 72 h co-culture with indicated MOIs of OV19t.

To visually assess the ability of the OV to infect and generate surface expression of CD19t on tumor cells, we infected the human triple-negative breast cancer cell line, MDA-MB-468, with OV19t for 16 h at varying multiplicity of infection (MOI), and stained tumor cells to detect expression of intracellular vaccinia virus and cell surface CD19t. We observed intracellular OV19t as well as surface expression of CD19t in an MOI-dependent manner (Fig. 1B). Interestingly, the intensity of CD19t expression following viral infection appeared higher than with MDA-MB-468 cells that were lentivirally transduced to stably express CD19t under a constitutive EF1α promoter. Expression of CD19t on tumor cells was evaluated at varying MOIs (0.00625 – 1) by flow cytometry at 48 h (Fig. 1C), and further quantified in a time-course from 24 to 72 h (Fig. 1D). Importantly, we observed nearly 100% CD19t expression on tumors cells after 24 h at an MOI 1, and after 72 h at lower MOIs even though 70% of tumor cells remained viable at that timepoint, providing a window of opportunity for targeting by CD19-specific CAR T cells. Similar trends of infection efficiency, CD19 expression, and OV-mediated killing of tumor cells were observed across multiple tumor types, including Capan-1, DU145, OV90, Panc-1, UM-SCC-1, UM-SCC-47, and U251T cells (Fig. 1E).

### OV delivery of CD19t to solid tumor cells redirect activity and cytotoxicity of CD19-CAR T cells in vitro

We next assessed whether hCD19t delivered to tumors by OV could activate CD19-CAR T cells. Tumor cells were infected with OV19t at varying MOIs and co-cultured with CD19-CAR T cells at an effector:target (E:T) ratio of 1:2 for 24 h. Cell surface expression of CD25 and 4-1BB (CD137) were used as markers of T cell activation and quantified by flow cytometry. At 24 h, CD19-CAR T cells demonstrated activation against OV19t-infected MDA-MB-468 cells in an MOI-dependent manner (Fig. S2, A and B). CD19-CAR T cells were also activated against OV19t-infected MDA-MB-231BR (brain-seeking human triple-negative breast cancer) cells after 24 h with similar kinetics as seen with MDA-MB-468 cells (Fig. S3A). CD19-CAR T cell function was then evaluated by measuring intracellular IFNγ levels and cell-surface CD107a as an early measure of effector activity following 16 h incubation with MDA-MB-468 cells infected with OV19t. CD19-CAR T cells showed robust activity when co-cultured with tumor cells infected with OV19t in an MOI-dependent manner (Fig. 2A).

**Fig. 2.**
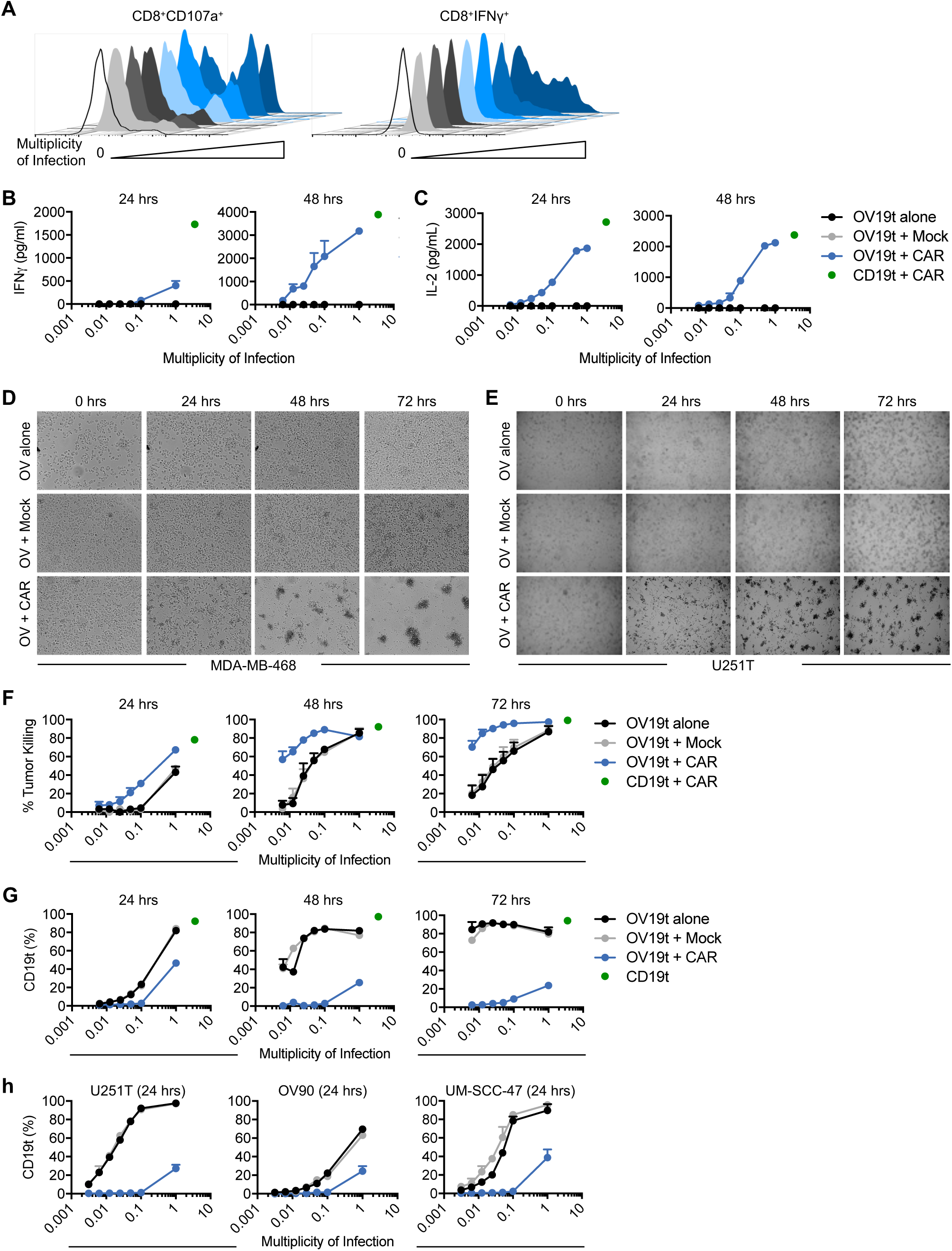
OV19t introduce expression of CD19t in tumor cells, which re-direct activation and cytotoxicity of CD19-CAR T cells *in vitro*. A) Representative flow cytometric analysis showing cell surface CD107a (left) and intracellular IFNγ expression (right) in CD8^+^CAR^+^ T cells following a 16 h co-culture with MDA-MB-468 tumor cells at a 1:1 effector:tumor (E:T) ratio with or without treatment with indicated MOI of OV19t. B) IFNγ and c) IL-2 production measured by ELISA in supernatants collected from co-cultures with or without treatment with OV19t at indicated MOIs for 24 and 48 h. D, E) Tumor killing assay of MDA-MB-468 (D) and U251T (E) visualized by phase-contrast microscopy at 10X magnification, and f) assessed by flow cytometry comparing Mock or CD19-CAR T cells following a 24, 48, or 72 h co-culture with MDA-MB-468 tumor cells treated with indicated MOIs of OV19t. G) CD19t expression on tumor cells in killing assay described in F). Killing data are presented as duplicates from two combined experiments. H) CD19t expression on U251T, OV90, and UM-SCC-47 cells treated with indicated MOIs of OV19t and co-cultured for 24 h with Mock or CD19-CAR T cells. Green dots indicate MDA-MB-468 cells stably expressing CD19t co-cultured with (B, C, F) or without T cells (G).

To further quantify the CD19-CAR T cell activity against tumor cells infected by OV19t, supernatants from co-cultures of CD19-CAR T cell and tumor cells with OV19t at varying MOI were collected to evaluate CAR-dependent cytokine production. At 24 h, IFNγ secretion was only observed at an MOI of 1. However, at 48 h, IFNγ secretion was detected in an MOI-dependent manner, reaching levels nearly equivalent to CD19-CAR T cells co-cultured with the positive control, MDA-MB-468 lentivirally transduced to stably express CD19t (Fig. 2B). Similar trends were observed with IL-2 secretion at each time point (Fig. 2C). To evaluate whether CD19-CAR activity leads to killing of OV19t-infected tumor cells, we performed *in vitro* tumor killing assays with MDA-MB-468 and U251T and qualitatively evaluated by phase-contrast microscopy in a 72 h time course with OV19t at an MOI of 0.05 (Fig. 2, D and E). We observed significantly greater killing of tumor cells infected with OV19t and co-cultured with CD19-CAR T cells at all time points when compared with OV19t-infected tumors alone. We further quantified killing ability against MDA-MB-468 cells using flow cytometry. As shown in Fig. 2F, we observed better killing of tumor cells infected with OV19t and co-cultured with CD19-CAR T cells at 24 h. At 48 and 72 h, the lowest MOI of OV19t (0.00625) induced suboptimal virus-mediated lysis of tumor cells (0 – 20%), whereas the combination of OV19t with CD19-CAR T cells showed potent tumor killing (60 – 70%). Similar combination effects were observed with MDA-MB-231BR cells (Fig. S3B). Importantly, we show CAR T cell-selective tumor killing highlighted by a dramatic reduction in CD19t-positive tumor cells in the combination group at all indicated time points and MOIs evaluated (Fig. 2G). Comparable trends in reduction of CD19t-positive tumor cells were observed using U251T, OV90, and UM-SCC-47 cells (Fig. 2H). We observed no CD19t expression or enhancement in tumor killing by CAR T cells when tumor cells were infected with OV-*tk* lacking CD19t (Fig. S4, A and B). Taken together, our data suggest that OV are capable of delivering a *de novo* tumor antigen, in this case CD19, and inducing antigen-specific CAR T cell-mediated anti-tumor activity.

### Anti-tumor efficacy of combination therapy of OV19t and CD19-CAR T cells in human tumor xenograft models

To first evaluate the *in vivo* anti-tumor activity of OV19t and the dynamics of CD19t expression in infected tumors, we treated mice bearing subcutaneous human MDA-MB-468 triple-negative breast tumors with a single intratumoral (i.t.) injection of OV19t. Choi *et al.* recently showed significant inhibition of tumor growth following i.t. treatment in MDA-MB-468-bearing nude mice ^10^. Therefore, we used varying doses [10^5^, 10^6^, 10^7^ plaque forming units (pfu) per mouse] to determine the infection efficiency and CD19t expression in tumors at 3, 7, and 10 d by flow cytometry (Fig. 3A). We determined that 10^7^ pfu of OV19t per mouse was optimal for combination with CD19-CAR T cells on day 10 post-infection in this model, showing approximately 70% of tumor cells expressing CD19t. We then performed an *in vivo* combination study with i.t. delivery of OV19t at 10^7^ pfu per mouse, followed 10 d later by i.t. treatment with 5 x 10^6^ of either Mock (untransduced) or CD19-CAR T cells. As expected, MDA-MB-468-bearing mice treated with Mock or CD19-CAR T cells alone showed no anti-tumor activity. Mice treated with OV19t alone or OV19t in combination with Mock T cells slowed tumor growth. Importantly, the combination of OV19t and CD19-CAR T cells exhibited a marked tumor regression (Fig. 3, B and C). We also confirmed the therapeutic benefits of this combination approach using U251T glioblastoma tumor-bearing mice (Fig. 3, D and E).

**Fig. 3.**
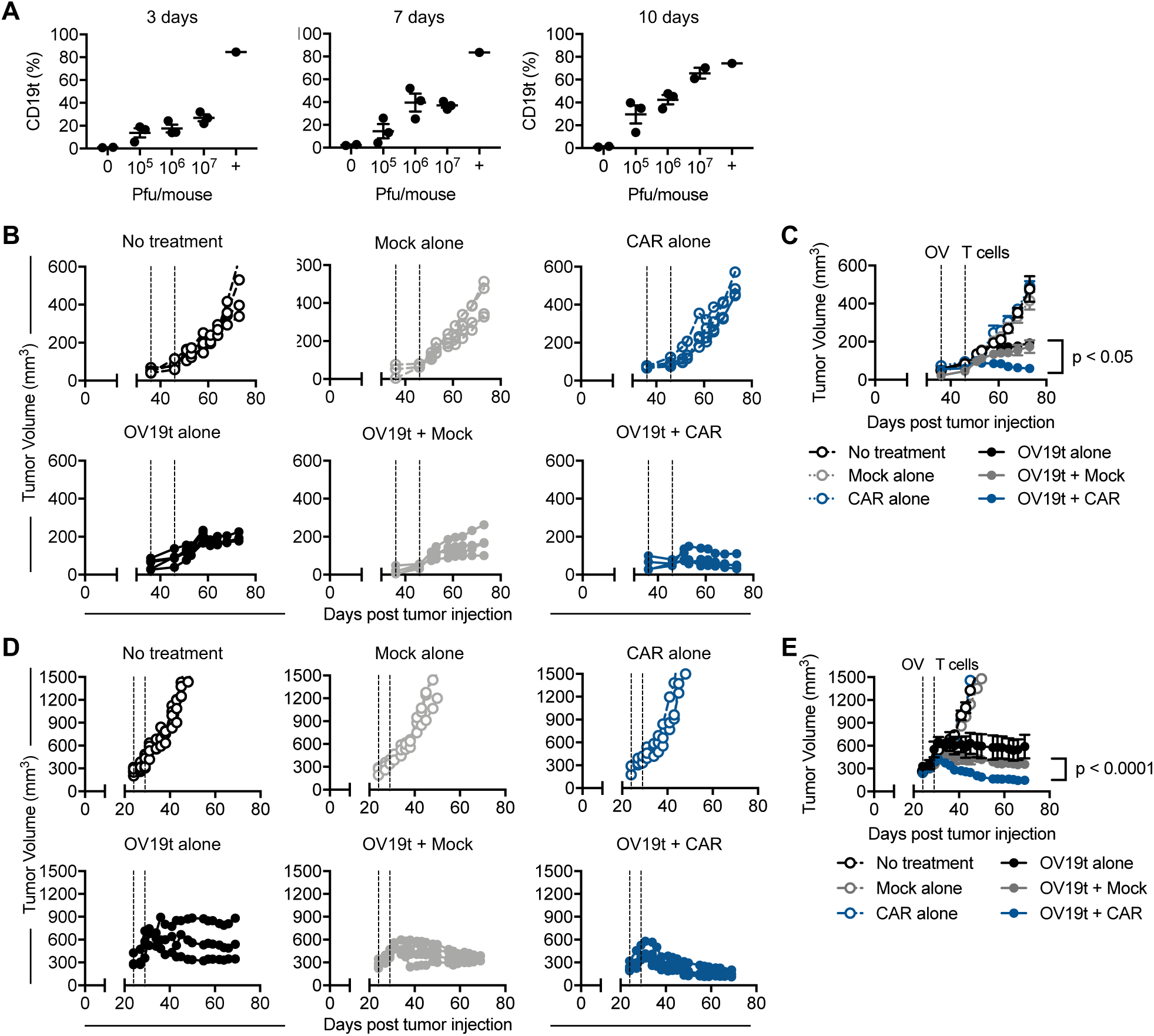
Anti-tumor efficacy of combination therapy of OV19t and CD19-CAR T cells in human xenograft tumor models. A) Mice were engrafted with subcutaneous MDA-MB-468 tumors (5×10^6^ cells) and at day 24 mice were intratumorally treated with 0, 10^5^, 10^6^, or 10^7^ plaque-forming units (pfu) per mouse, which were harvested at day 3 (left), 7 (middle), or 10 (right) after treatment for quantification of CD19t expression via flow cytometry. The positive control (+) represents MDA-MB-468 tumors previously lentivirally transduced to stably express CD19t. B, C) Tumor volume (mm^3^) in NSG mice bearing subcutaneous MDA-MB-468 (5×10^6^ cells) tumors on day 0, and treated with OV19t (10^7^ pfu) on day 36. On day 46, mice were treated with either Mock or CD19-CAR T cells (5×10^6^ cells). Lines in (B) represent individual mice per group, which are averaged in (C). n ≥ 4 per group. Data presented in (C) are mean ± standard error mean. All data are representative of two independent experiments. D, E) Tumor volume (mm^3^) in NSG mice bearing subcutaneous U251T (5×10^6^ cells) tumors on day 0, and treated with OV19t (10^3^ pfu) on day 24. On day 29, mice were treated with either Mock or CD19-CAR T cells (5×10^6^ cells). Lines in (D) represent individual mice per group, which are averaged in (E). n ≥ 3 per group. Data presented in (E) are mean ± standard error mean.

### Anti-tumor efficacy of combination therapy of OVm19t and CD19-CAR T cells in a murine immunocompetent tumor model

We next evaluated the therapeutic benefits of this combination approach in an immunocompetent mouse tumor model. To that end, we generated OV19t with the murine CD19t gene (OVm19t), and utilized murine splenic T cells to manufacture CD19-CAR T cells. The murine CD19-CAR was generated with a retrovirus containing the anti-mouse CD19 single-chain variable fragment (scFv) antigen-binding domain derived from the 1D3 hybridoma as previously described^11^, a murine CD8 hinge extracellular spacer and transmembrane domain, a 4-1BB intracellular signaling domain, and a CD3. The truncated human epidermal growth factor receptor (EGFRt) for cell tracking was separated from the CAR with a T2A skip sequence (Fig. S5A). Murine CD19-CAR T cells potently killed MC38 cells lentivirally engineered to express murine CD19t (Fig. S5B). To evaluate whether murine CD19-CAR T cell activity leads to killing of OVm19t-infected MC38 cells, we performed *in vitro* tumor killing assays and quantified killing ability using flow cytometry. As shown in Fig. 4A (left), we observed better killing of tumor cells infected with OVm19t and co-cultured with CD19-CAR T cells at 24 h compared with OVmCD19t alone. We observed intracellular OVm19t as well as surface expression of CD19t in an MOI-dependent manner, and CAR T cell-selective tumor killing highlighted by a dramatic reduction in CD19t-positive and OVm19t-positive tumor cells in the combination group at all MOIs evaluated (Fig. 4A, middle and right). These data suggest a similar combination anti-tumor effect of OV carrying CD19t and CD19-CAR T cells using a fully murine system.

**Fig. 4.**
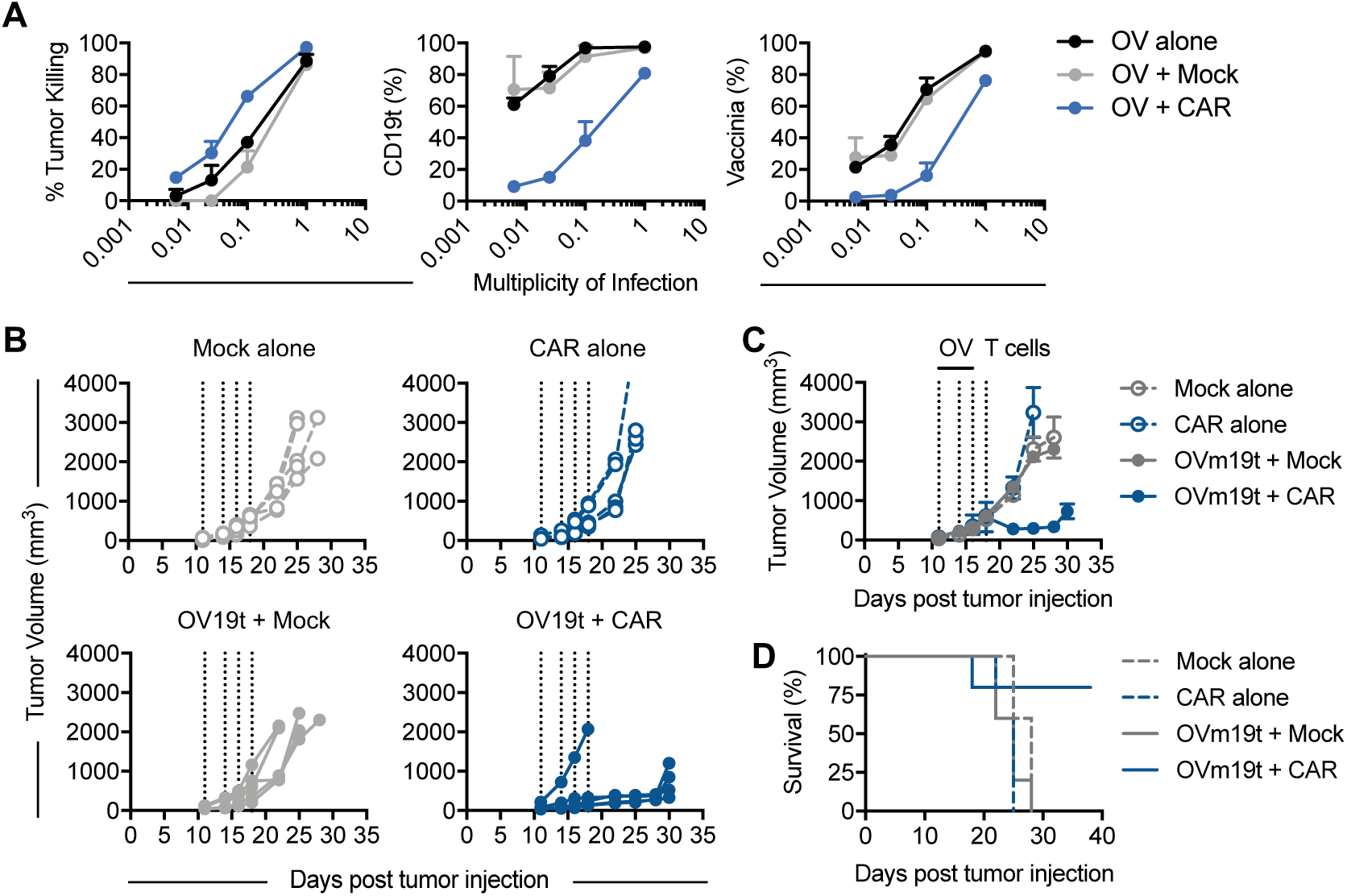
Anti-tumor efficacy of combination therapy of OVm19t and CD19-CAR T cells in an immunocompetent murine syngeneic tumor model. A) Tumor killing assay assessed by flow cytometry comparing Mock or CD19-CAR T cells following a 24 h co-culture with MC38 tumor cells treated with indicated MOIs of OVm19t (left graph). Quantification of CD19t (middle graph) and vaccinia (right graph) expression on tumor cells in the same tumor killing assay. Mice were engrafted with subcutaneous MC38 tumors (1×10^6^ cells) and at day 11, 14, and 16 mice were intratumorally treated with 0 or 10^7^ plaque-forming units (pfu) per mouse. On day 18, mice were treated with either Mock or CD19-CAR murine T cells (5×10^6^ cells). Tumor volume (mm^3^) was measured by calipers. Lines represent individual mice per group, which are averaged in (C). n = 4 per group. Data presented in (C) are mean ± standard error mean. D) Kaplan-Meier survival curves from experiment in (B). All data are representative of two independent experiments.

The *in vivo* anti-tumor activity of this combination approach in a syngeneic tumor model was tested using MC38 tumor-bearing C57BL/6j mice. Minimal anti-tumor activity was observed using Mock (untransduced) or CD19-CAR T cells alone in this tumor model. While three doses of 10^7^ pfu of OVm19t injected intratumorally every other day also demonstrated minimal anti-tumor effects, we observed a significant delay in tumor growth in mice treated with OVm19t in combination with CD19-CAR T cells (Fig. 4, B and C), with prolonged survival of mice in the combination group (Fig. 4D). Taken together, we demonstrate the broad applicability of our combination immunotherapy approach with OV-mediated delivery of CD19t antigen and CD19-CAR T cells using both human xenograft and immunocompetent mouse tumor models.

### CD19-CAR T cell-mediated tumor killing promotes the early release of OV19t

One of the potential pitfalls of using OV to deliver CAR T cell target antigens to tumor cells is the suboptimal or non-uniform infection of solid tumors^12,13^. To address this, we assessed whether CAR T cell-mediated killing of virally-infected tumor cells could enhance the release of oncolytic viral particles for subsequent infection of surrounding tumor cells. Tumor cells were first infected with varying MOI of OV19t and then incubated with Mock or CD19-CAR T cells. Viral titer was then measured in collected supernatants. As shown in Fig. 5A, the combination of OV19t and CD19-CAR T cells demonstrated significant increases in released viral particles, compared with OV19t infection alone or the combination with Mock T cells. We further demonstrated that viral supernatants showed increased infection of fresh tumor cells (Fig. 5, B and C), which resulted in significantly greater tumor killing potential (Fig. 5, D and E) due to increased release of OV when combined with CD19-CAR T cells. The release of oncolytic viral particles from dying tumor cells by antigen-specific CD19-CAR T cells suggests that we may enable more homogeneous infection of tumor cells by this novel combination therapy.

**Fig. 5.**
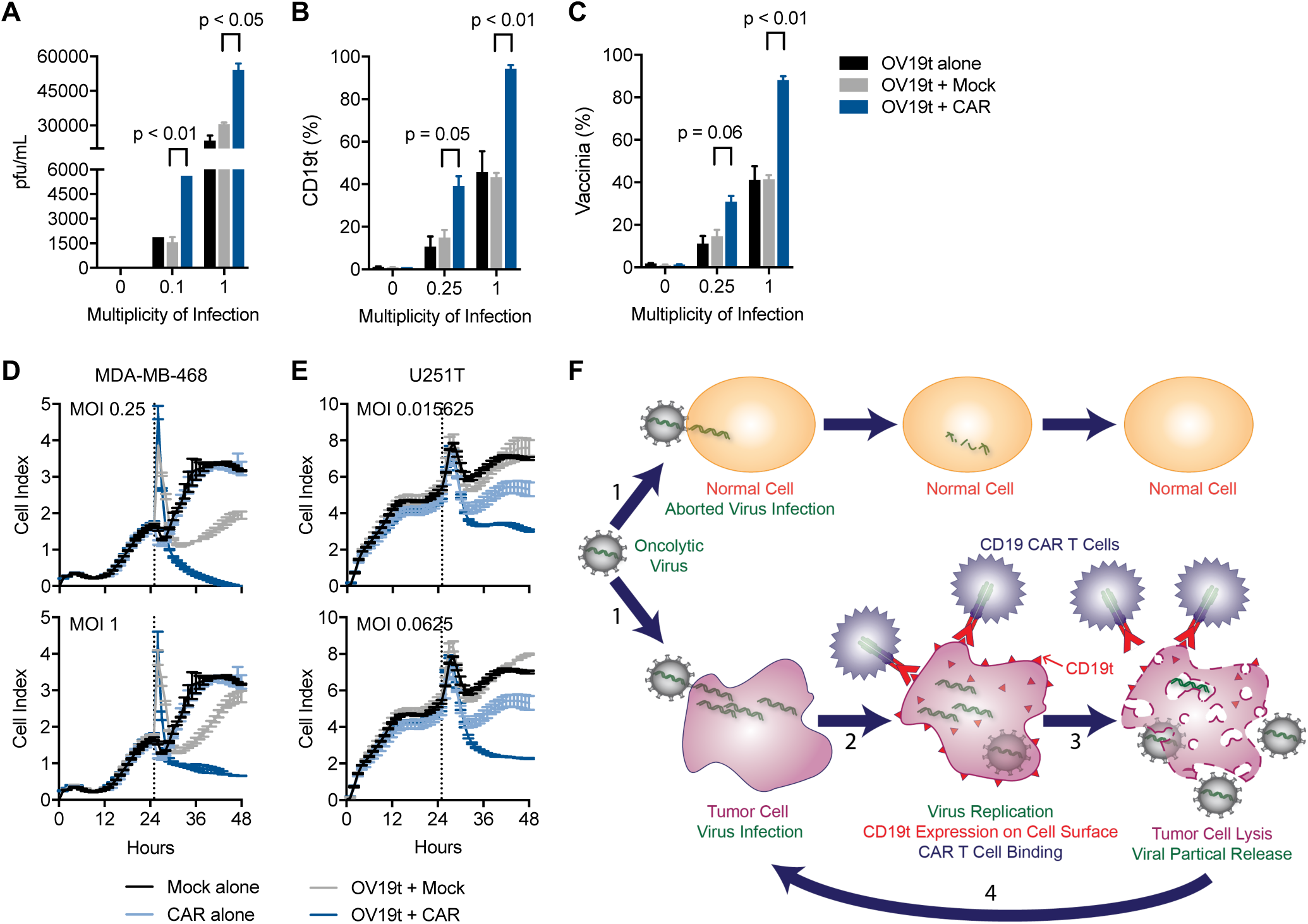
CD19-CAR T cell-mediated tumor killing promotes the early viral particle release and infection of tumor cells. A) OV19t at varying MOIs were added to MDA-MB-468 tumor cells for 16 h and then incubated with Mock or CD19-CAR T cells for an additional 4 h. Viral titer was then measured in collected supernatants. B, C) OV19t at varying MOIs were added to MDA-MB-468 tumor cells for an initial 3 h incubation. Virus was washed off, and cells were incubated for an additional 4 h prior to adding Mock or CD19-CAR T cells for 18 h. Supernatants were collected and added to freshly plated MDA-MB-468 tumor cells and incubated for 18 h prior to analysis by flow cytometry for expression of CD19t (B) and vaccinia (C). A similar experimental design was performed (as in B and C), and MDA-MB-468 (D) and U251T (E) were monitored using an xCELLigence RTCA system. F) Schematic of combination therapy concept utilizing OV to introduce CAR targets to solid tumors: 1) Virus infection of normal or tumor cells using OV19t. 2) Virus replication and CD19t expression on cell surface, allowing for CD19-CAR T cell targeting. 3) Tumor cell lysis leading to viral particle release. 4) Released viral particles re-initiate virus infection of surrounding tumor cells.

## Discussion

We believe that our findings address a major challenge facing CAR T cell therapies for solid tumors, which include targeting surface proteins that demonstrate heterogeneous expression patterns in tumors, and shared expression of these surface proteins on some normal tissues. We have demonstrated the distinct capability of OV to selectively deliver a CAR-targetable tumor antigen to a human xenograft model of triple-negative breast cancer. Importantly, the combination with CD19-CAR T cells in this model resulted in greater viral spread and the potential for more uniform expression of the CAR target in the tumor. This antigen delivery method may be adopted in other solid cancer types that lack amenable tumor antigens for safe and effective targeting by CAR T cells. CAR T cell therapy and oncolytic virotherapy are complementary modalities for cancer treatment with remarkable potential. While the introduction of *de novo* tumor targets distinguishes our approach from other recent studies combining OV with CAR T cells^14,15^, we plan to further modify these OV to express checkpoint pathway inhibitors, cytokines, and chemokines to augment CAR T cell trafficking and anti-tumor activities.

While we performed these proof-of-concept studies using CD19t as a target, this approach has the potential to introduce other targets, including non-human species-specific antigens^16^, which may also further enhance the anti-tumor efficacy by eliciting host immunity to foreign proteins. However, targeting of CD19 with CAR T cells is a clinically-validated approach for treating hematological malignancies, and has demonstrated clinically-manageable B-cell aplasia^5,6^, supporting the future clinical development in this potentially safe and effective combination strategy. Also, limited persistence of vaccinia viruses is in part due to antibody-mediated clearance of virus particles^17^, which may be significantly dampened by depleting B cells with CD19-CAR T cells, thus improving systemic and local persistence of OV19t.

Host immunity is important to the outcome of certain OV, as has been shown for HSV^18^. Additionally, T and NK cells have been identified as major components of host immunity after vaccinia infection and contribute to OV-mediated anti-tumor effects^19^. One might envision that following the combination of OV19t and CD19-CAR T cells, new tumor antigens released from dying tumor cells may instigate another wave of anti-tumor immunity mediated by adoptively transferred or endogenous T cells. While we demonstrated robust anti-tumor activity of our combination approach in an immunocompetent mouse tumor model, future studies with this model will not only enable studying the global effects of the OV alone and in combination with CAR T cells, but also enable us to better understand the changes that occur locally in the tumor microenvironment, as well as systemically, following combination therapy. Further, we envision that other oncolytic viruses besides vaccinia (e.g. HSV, Maraba virus, and adenoviruses) can also be exploited in this combination strategy, and that the immunogenic aspects of the therapy may vary depending on the virus.

In summary, we have demonstrated that OV can effectively deliver the CD19-CAR target to solid tumors, which are then susceptible to CD19-CAR T cell-mediated tumor destruction, and further CAR T cell-mediated viral spread (Fig. 5F). These findings potentially extend the use of clinically-approved CD19-CAR T cells beyond B-cell malignancies and into the treatment paradigm for multiple solid tumors, warranting further investigations of OV19t, or similar viruses delivering CAR targets, in combination with CAR T cells for the treatment of intractable solid tumors.

## Materials and methods

### Cell lines and viruses

Human triple-negative breast cancer cell line MDA-MB-468 (ATCC HTB-132) was cultured in Dulbecco’s Modified Eagle Medium (DMEM) containing 10% fetal bovine serum (FBS, Hyclone) and 1x antibiotic antimycotic (AA, Gibco) supplemented with 25 mmol/L HEPES (Irvine Scientific) and 2 mmol/L L-glutamine (Thermo Fisher Scientific) (complete DMEM). The brain-seeking human triple-negative breast cancer cell line MDA-MB-231BR^20^ (a kind gift from Dr. Patricia S. Steeg, NIH, Bethesda, MD) was cultured in DMEM/Ham F-12 (F12; 1:1) containing 10% FBS and 1x AA. Human pancreatic cancer cell line Capan-1 (ATCC HTB-79) was cultured in Iscove’s Modified Dulbecco’s Medium (IMDM) containing 20% FBS and 1x AA. Human pancreatic cancer cell line Panc-1 (ATCC CRL-1469) was cultured in Roswell Park Memorial Institute (RPMI) containing 10% FBS and 1x AA. Human ovarian cancer cell line OV90 (ATCC CRL-11732) was cultured in 1:1 volume of MCDB 105 medium (Sigma-Aldrich) and Medium 199 (Gibco) containing 20% FBS and 1x AA. Human head and neck carcinoma line UM-SCC-1 (EMD Millipore) was cultured in DMEM containing 20% FBS, 1x AA, and 1x non-essential amino acids (NEAA, Life Technologies). Human head and neck carcinoma line UM-SCC-47 (EMD Millipore) was cultured in DMEM containing 10% FBS, 1x AA, and 1x NEAA. Human prostate cancer cell line DU145 (ATCC HTB-81) was cultured in RPMI containing 10% FBS and 1x AA. Human glioblastoma cell line U251T (gift from Dr. Waldemar Debinski, Wake Forest School of Medicine) was cultured in complete DMEM. Human embryonic kidney cell line 293T (ATCC CRL-3216) and human fibrosarcoma cell line HT1080 (ATCC CCL-121) were cultured in complete DMEM. African green monkey kidney fibroblasts (CV-1) (ATCC CCL-70) were cultured in DMEM containing 10% FBS and 1x AA. CV-1 cells were used for both amplification and titration of orthopoxviruses. Murine colon adenocarcinoma cell line MC38 was cultured in complete DMEM.

### Generation of recombinant chimeric orthopoxvirus expressing human and murine truncated CD19 (CD19t)

To generate a shuttle vector containing the human (hCD19t) and murine (mCD19t) CD19t expression cassette with the VACV synthetic early (SE) promoter, the hCD19t and mCD19t cDNAs were PCR-amplified from the plasmids hCD19t-2A-IL2-pHIV7 and mCD19t-epHIV7 using Q5 High-Fidelity 2X Master Mix (New England Biolabs Inc., Ipswich, MA) and the primers: 5’-GCG GTC GAC CAC CAT GCC ACC TCC TCG CCT CCT CTT CTT CCT CCT CTT CCTC-3’ and 5’-GCG GGA TCC ATA AAA ATT AAT TAA TCA TCT TTT CCT CCT CAG GAC CAG GGC TCT TTG AAG ATG-3’. The PCR fragment was digested with *Sal* I and *Bam*H I and cloned into the same-cut p33NCTK-SE-hNIS replacing hNIS to yield p33NCTK-SE-hCD19t and p33NCTK-SE-mCD19t. The hCD19t and mCD19t cDNAs in p33NCTK-SE-hCD19t and p33NCTK-SE-mCD19t were confirmed by sequencing. CV-1 cells were infected with CF33^10^ at a multiplicity of infection (MOI) of 0.1 for 1 h and then transfected with p33NCTK-SE-hCD19t and p33NCTK-SE-mCD19t by use of jetPRIME *in vitro* DNA & siRNA transfection reagent (Polyplus-transfection Inc., New York, NY). Two days post infection, infected/transfected cells were harvested and the recombinant viruses (OV19t) were selected and plaque purified as described previously^21^.

### DNA constructs

MDA-MB-468 and MDA-MB-231BR cells were engineered to express human CD19t by transduction with epHIV7 lentivirus carrying the human CD19t gene under the control of the EF1α promoter. A similar procedure was used to engineer MC38 cells to express murine CD19t. The human CD19-28ζ CAR lentiviral construct with truncated human EGFR (EGFRt) separated by a T2A ribosome skip sequence was previously described^22^. The murine CD19-BBζ CAR retroviral construct (pMYs, Cell Biolabs Inc.) contained an anti-mouse CD19-targeted single-chain variable fragment (scFv) sequence derived from the 1D3 hybridoma^11^, a murine CD8 hinge extracellular spacer and transmembrane domain, a murine 4-1BB intracellular costimulatory signaling domain, a murine CD3ζ cytolytic domain, and the human EGFRt separated from the CAR by a T2A ribosome skip sequence.

### Human T cell enrichment, lentivirus production and transduction, and ex vivo expansion

T-cell isolation, lentivirus production and transduction, and *ex vivo* expansion of CAR T cells was performed as previously described^23^.

### Murine retrovirus production

Retrovirus was generated by plating PlatE cells for 1 week in complete DMEM with the addition of selection antibiotics including 1 µg/mL puromycin (InvivoGen) and 10 µg/mL blasticidin (InvivoGen). One day prior to production, cells were washed once and cultured with warm antibiotic-free complete DMEM. On the day of transfection, cells were washed again and 6 µg of plasmid DNA was added with FugeneHD transfection reagent (Promega). Supernatants were collected at 24, 36, and 48 h and frozen at -80°C until further use.

### Murine T cell enrichment, transduction, and ex vivo expansion

Mouse splenocytes were obtained from naïve C57BL/6j mice (The Jackson Laboratory, Bar Harbor, Maine, USA), and T cells were isolated using the EasySep Mouse T Cell Isolation Kit (StemCell Technologies) according to the manufacturer’s protocol. Freshly isolated mouse T cells were cultured in RPMI media containing 10% FBS (Hyclone), 50 U/mL recombinant human IL-2 (Novartis Oncology), 10 ng/mL recombinant murine IL-7 (Peprotech), and 50 µM 2-Mercaptoethanol (Gibco). For CAR retroviral transduction, T cells were cultured with mouse CD3/CD28 Dynabeads (Invitrogen) overnight, and plated onto Retronectin (Takara) coated 24-well non-treated tissue culture plates (Corning Life Sciences) with 1 mL of murine CD19-CAR retrovirus. Plates were spinoculated at 1500 g for 1 h at 32°C. After culturing the transduced cells for 4 days, beads were magnetically removed and T cells were used for *in vitro* functional assays and *in vivo* tumor models. Purity and phenotype of CAR T cells were confirmed by flow cytometry.

### Intracellular/extracellular staining and flow cytometry

Flow cytometric analysis was performed as previously described^23^. Tumor cells were identified using Ep-CAM (CD326) (BioLegend, Clone: 9C4). Cell viability was determined using LIVE/DEAD Fixable Violet Dead Cell Stain Kit for 405 nm excitation (Invitrogen). For vaccinia staining, cells were fixed and permeabilized using BD Cytofix/Cytoperm Fixation/Permeabilization Solution Kit according to the manufacturer’s protocol (BD Biosciences). Cells were then incubated with anti-vaccinia virus primary antibody (Abcam) and incubated for 30 minutes at 4°C in the dark. The cells were washed twice prior to secondary stain with Goat Anti-Rabbit IgG H&L (Alexa Fluor 488) (Abcam) for 30 minutes at 4°C in the dark. The cells were then washed twice prior to resuspension in FACS buffer and acquisition on the MACSQuant Analyzer 10 (Miltenyi Biotec). Data were analyzed with FlowJo software (v10, TreeStar).

### OV transduction and T-cell functional assays

For OV transduction and tumor killing assays, CAR T cells and tumor targets were co-cultured at varying E:T ratios along with the addition of varying MOIs of OV19t in complete X-VIVO in the absence of exogenous cytokines in round-bottom 96-well tissue culture-treated plates (Corning) for 1-3 d and analyzed by flow cytometry as described above. Tumor killing by CAR T cells with or without OV19t was calculated by comparing CD45-negative cell counts relative to that observed by Mock (untransduced) T cells from the same healthy donor. For T cell activation assays, CAR T cells and tumor targets were co-cultured at an E:T ratio of 1:2 along with the addition of varying MOIs of OV19t in complete X-VIVO in the absence of exogenous cytokines in 96-well plates for 1-3 d and analyzed by flow cytometry for specific markers of T cell activation. A similar procedure was followed for mouse functional assays using OVmCD19t, using complete RPMI instead of X-VIVO, and at an E:T ratio of 1:1. For degranulation and intracellular cytokine assays, CAR T cells and tumor targets were co-cultured at varying E:T ratios along with the addition of varying MOIs of OV19t in complete X-VIVO in the absence of exogenous cytokines in round-bottom 96-well plates. FITC-CD107a was added to the cultures for 16-18 h at 37°C, and then cells were fixed and permeabilized before analysis by flow cytometry.

### Viral Plaque Assay

Following infection of cells with virus as mentioned above, supernatants were collected and processed as previously described^10^.

### Immunofluorescence Microscopy

Tumor cell lines were analyzed for vaccinia and CD19t expression via immunofluorescence microscopy. 4 × 10^4^ MDA-MB-468 and MDA-MB-468-CD19t cells were plated in individual wells of 96-well glass bottom plates (Cellvis), and incubated for 24 h to allow for adherence. Media was aspirated from each well and gently washed with PBS + 1% BSA. Cells were stained with primary CD19 antibody (Abcam) at 4°C for 1 h, washed, and stained with secondary streptavidin AlexaFluor 647 (Invitrogen) for 1 h at RT. Cells were then fixed and permeabilized using Cytofix/Cytoperm Fixation/Permeabilization Solution (BD Biosciences) for 10 min at RT. The solution was aspirated and stained with anti-vaccinia antibody (Abcam) for 1 h at RT, washed, and stained with goat anti-rabbit IgG AlexaFluor 488 (Abcam) for 1 h. Cells were stained with DAPI for 5 minutes at RT and each well was imaged at 10-63X magnification.

### Cytokine ELISA

Tumor cells and CD19-CAR T cells were plated into 96-well round bottom plates (Costar) and varying MOIs of OV19t (OVmCD19t) were added. Following incubations at 37°C for 24, 48, or 72 h, supernatants were collected and run according to the Human or Mouse IFNγ or IL-2 ELISA Ready-SET-GO! (eBioscience) manufacturer’s protocol. Plates were read at 450 nm using the Cytation 3 Cell Imaging Multi-Mode Reader and Gen5 Microplate Reader and Imager Software (BioTek).

### xCELLigence

The xCELLigence real-time cell analysis (RTCA) instrument (ACEA Biosciences) was used for impedance experiements to determine tumor cell killing according to the manufacturer’s protocol. Briefly, tumor cells were plated at 25,000 cells per well and co-cultured with varying MOIs of OV19t and incubated at 37°C for 3 h. Fresh media was added to remove any residual viral particles. Next, the cells were incubated for an additional 4 h, and Mock or CAR T cells were added for 18 h. Supernatants were collected and sonicated three times, and then added to freshly plated tumor cells to evaluate cell killing.

### In vivo tumor studies

All animal experiments were performed under protocols approved by the City of Hope Institutional Animal Care and Use Committee. For human tumor xenograft studies, MDA-MB-468 and MDA-MB-468-CD19t cells (5 × 10^6^ cells/mouse), or U251T cells (1 × 10^6^ cells/mouse), were prepared in HBSS^−/−^ and injected subcutaneously in the flank of female NSG mice. Tumor growth was monitored 2-3 times per week via caliper measurement. Once tumor volumes reached approximately 100-500 mm^3^, OV19t viruses prepared and diluted in PBS were intratumorally (i.t.) administered at 10^3^-10^7^ pfu/mouse. For combination therapy studies, CAR T cells were prepared in PBS and injected i.t. at either 5 or 10 d after OV treatment. For OV19t transduction studies, mice were euthanized and tumors were harvested and processed for flow cytometry (described above) at 3, 7, or 10 d post OV19t treatment. For all studies, mice were euthanized and tumors were harvested and processed for flow cytometry once tumors no greater than 15 mm in diameter.

For immunocompetent mouse studies, MC38 cells (5 × 10^5^ cells/mouse) were prepared in HBSS^−/−^ containing 1:1 v:v of growth-factor reduced matrigel (Corning Life Sciences) and injected subcutaneously in the flank of female C57BL/6j mice. Tumor growth was monitored 2-3 times per week via caliper measurement. Once tumor volumes reached approximately 100-300 mm^3^, OVm19t virus prepared and diluted in PBS was intratumorally (i.t.) administered for a total of 3 doses of 10^7^ pfu/mouse per dose. For combination therapy studies, CD19-CAR T cells were prepared in PBS and injected i.t. at 2 d after the last OV treatment.

### Statistical analysis

Data are presented as mean ± standard deviation (SD), unless otherwise stated. Statistical comparisons between groups were performed using the unpaired two-tailed Student’s t-test to calculate p-value, unless otherwise stated.

---

**Supplemental Figure 1:**
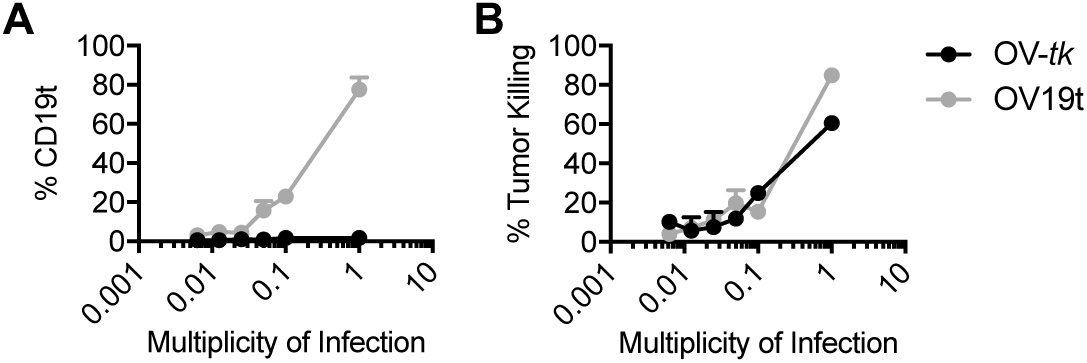
Replacing *tk* gene with hCD19t in OV does not significantly impact the infection efficiency or killing of MDA-MB-468. A) Quantification of cell surface CD19t expression following 24 h co-culture with indicated MOIs of OV-*tk* (control) or OV19t. B) Tumor killing assessed by flow cytometry comparing MDA-MB-468 treated with indicated MOIs of OV-*tk* (control) or OV19t.

**Supplemental Figure 2:**
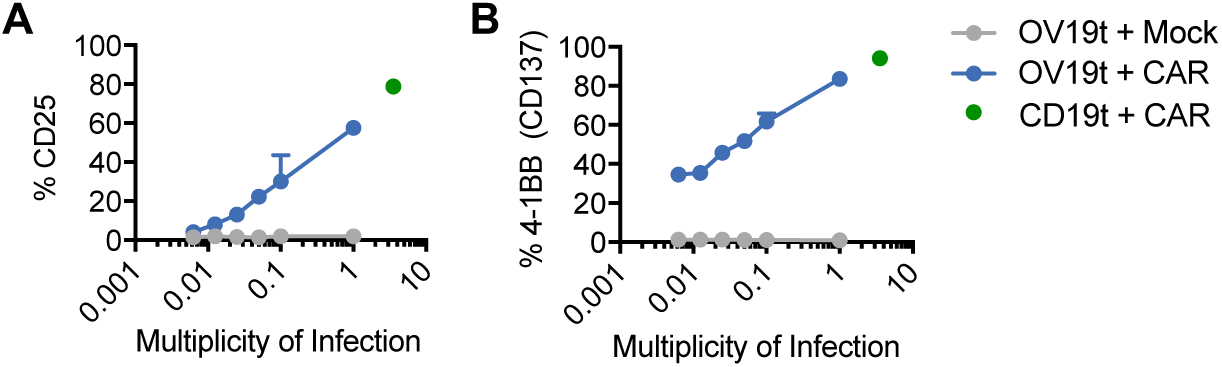
Activation of CD19-CAR T cells against OV19t infected tumors expressing CD19t. Quantification of A) CD25 and B) CD137 expression on Mock (untransduced) or CD19-CAR T cells following a 24 h co-culture with tumor cells at an E:T ratio of 1:2 with or without treatment with indicated MOI of OV19t. Green dots indicate T cells co-cultured with MDA-MB-468 cells stably expressing CD19t.

**Supplemental Figure 3:**
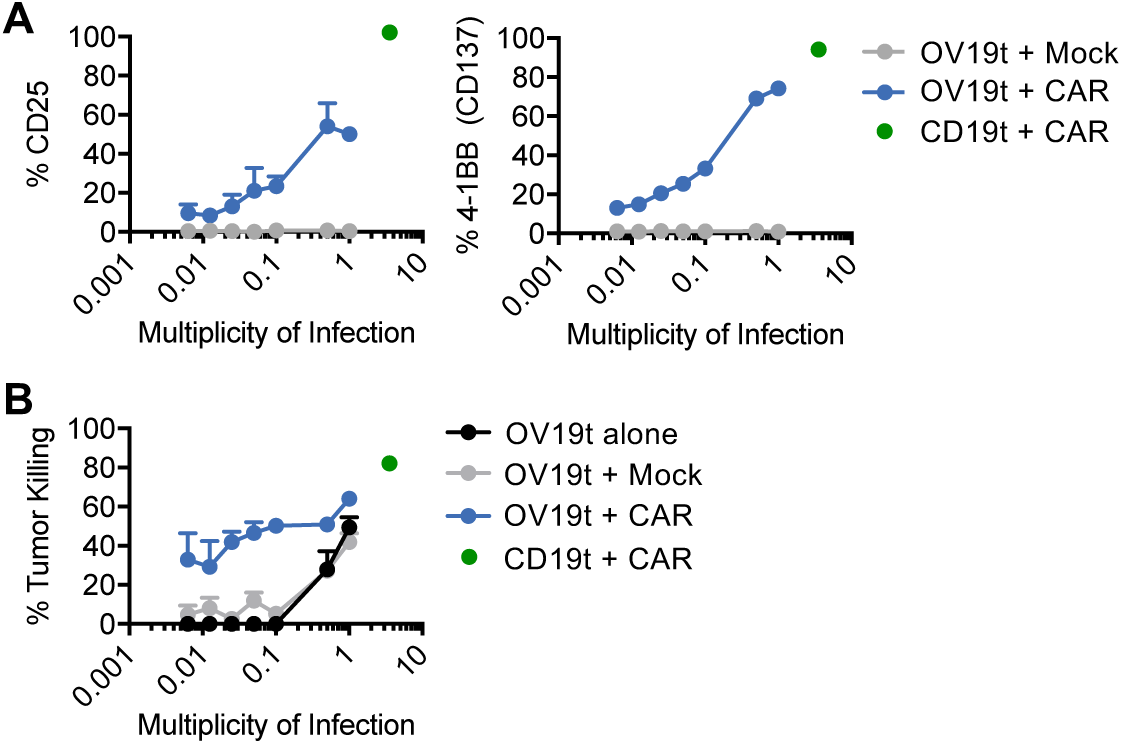
CD19-CAR T cells activate and kill MDA-MB-231 cells infected with OV19t. Quantification of CD25 (left) and CD137 (right) expression on Mock or CD19-CAR T cells following a 24 h co-culture with MDA-MB-231 cells at an E:T ratio of 1:2 with or without treatment with indicated MOI of OV19t. B) Tumor killing assessed by flow cytometry comparing Mock or CD19-CAR T cells following 24 h co-culture with MDA-MB-231 cells treated with indicated MOIs of OV19t. Green dots indicate T cells co-cultured with MDA-MB-231 cells lentivirally transduced to stably express CD19t.

**Supplemental Figure 4:**
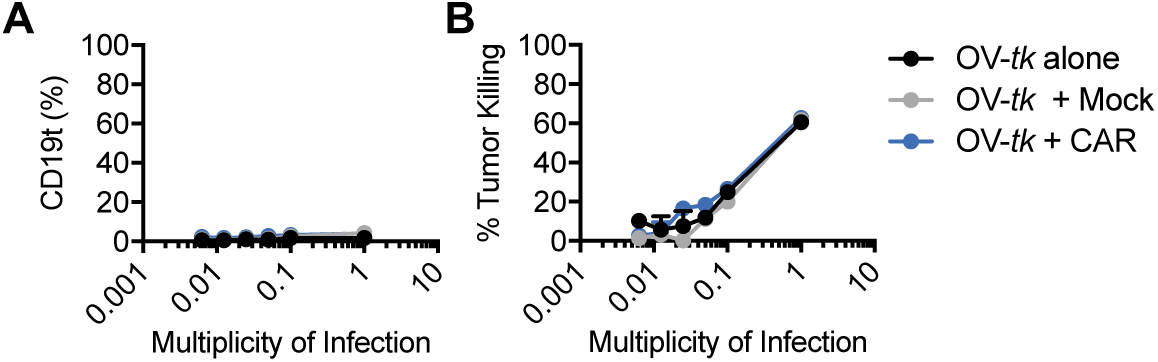
OV carrying *tk* does not induce CAR T cell activity. *A*) Quantification of cell surface CD19t expression following 24 h co-culture with indicated MOIs of OV-*tk* alone, or with Mock or CD19-CAR T cells. B) Tumor killing assessed by flow cytometry comparing MDA-MB-468 treated with indicated MOIs of OV-*tk* alone, or with Mock or CD19-CAR T cells.

**Supplemental Figure 5:**
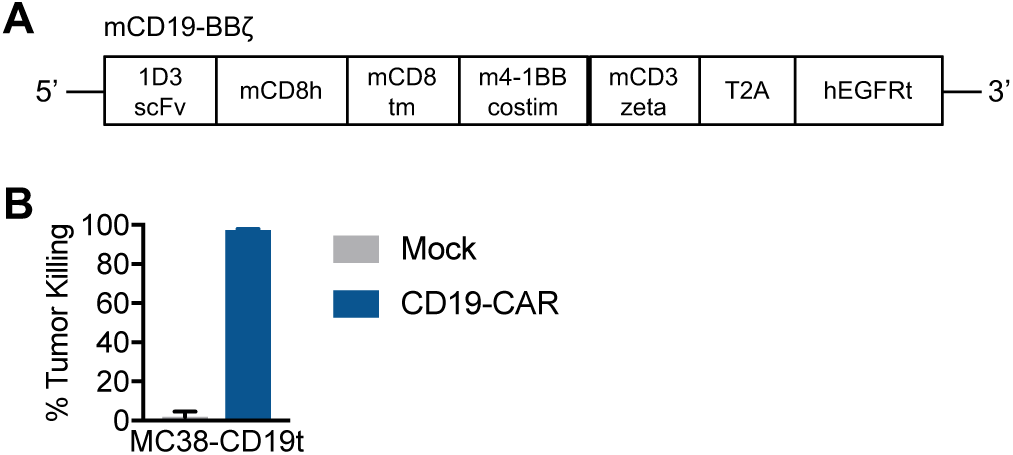
Murine CD19-CAR T cells. A) Diagram of the retroviral expression cassette with CD19-CARs containing the murine scFv (1D3 clone) targeting CD19, with a murine CD8 hinge and transmembrane domain, a murine cytoplasmic 4-1BB costimulatory domain, and a murine cytolytic CD3ζ domain. A truncated non-signaling EGFR (EGFRt), separated from the CAR with a T2A ribosomal skip sequence, was expressed for tracking CAR-expressing cells. B) Tumor killing assay (E:T ratio of 1:1) assessed by flow cytometry comparing Mock or CD19-CAR T cells following a 24 h co-culture with MC38 tumor cells lentivirally-transduced to stably express CD19t (MC38-CD19t). Tumor killing is compared to tumor cells cultured alone.

